# Meso-limbic Gene Expression Findings from Mouse Cocaine Self-Administration Recapitulate Human Cocaine Use Disorder

**DOI:** 10.1101/2020.01.31.929406

**Authors:** Spencer B. Huggett, Jason A. Bubier, Elissa J. Chesler, Rohan C. Palmer

## Abstract

Animal models of drug use have been employed for over 100 years to facilitate the identification of mechanisms governing human substance use and addiction. Most cross-species research on drug use/addiction examines behavioral overlap, but studies assessing neuro-molecular correspondence are lacking. Our study utilized transcriptome-wide data from the hippocampus and ventral tegmental area (VTA)/midbrain from a total of 35 human males with cocaine use disorder/controls and 49 male C57BL/6J cocaine/saline administering/exposed mice. We hypothesized that individual genes (differential expression) and systems of co-expressed genes (gene networks) would demonstrate appreciable overlap across mouse cocaine self-administration and human cocaine use disorder. We found modest, but significant associations between differentially expressed genes associated with cocaine self-administration (short access) and cocaine use disorder within meso-limbic circuitry, but non-robust associations with mouse models of acute cocaine exposure, (cocaine) context re-exposure and cocaine + context re-exposure. Investigating systems of co-expressed genes, we also found several validated gene networks with weak to moderate conservation between cocaine/saline self-administering mice and disordered cocaine users/controls. The most conserved hippocampal and VTA gene networks demonstrated substantial overlap (2,029 common genes) and included novel and previously implicated targets of cocaine use/addiction. Lastly, we conducted expression-based phenome-wide association studies of the nine *common* hub genes across conserved gene networks and found that they were associated with dopamine/serotonin function, cocaine self-administration and other relevant mouse traits. Overall, our study identified and characterized homologous transcriptional effects between mouse models of cocaine self-administration and human cocaine use disorder that may serve as a benchmark for future research.

## Introduction

For over a hundred years animal models of drug use have provided mechanistic insight into human drug addiction^1^. Most of the knowledge regarding the neurobiological mechanisms contributing to human drug addiction is derived from extrapolating findings from animal studies. The *modus operandi* for drug use in animals is the self-administration paradigm (18,865 publications between 2009-2019 via PubMed), in which, animals perform a behavior to consume a drug or saline. Numerous variations of drug self-administration exist to model addiction related behavior, but our study predominately focuses on the classic (short-access) cocaine self-administration paradigm, which is considered a valid model for reinforcement learning and voluntary cocaine use^2,3^. These features seem important for the development of substance abuse, but controversy remains pertaining to the relationship between classic animal self-administration paradigms and human drug addiction^4,5^.

Generally the translational relevance for animal models of drug use is investigated through a behavioral lens^6,7^, whereby researchers assess the overlay of *specific* animal behaviors with behavioral characteristics or clinical symptomology of human drug use disorder. Such an approach is based almost entirely on face validity of the animal model and the recapitulation of the highly complex aspects of disordered drug use. Transcriptomic methods provide an alternate means of assessing the construct validity of behavioral assays and their relevance to the study of the mechanisms of human conditions. But, despite an expanding body of bioinformatics data, studies investigating cross-species neuro-molecular overlap have been limited – begging the question of whether, or to what degree, gene expression profiles are comparable between animal models of substance use and human drug use disorder.

Human post-mortem brain studies have produced useful information regarding the molecular hallmarks associated with the “end point” of psychiatric disorders^8^. While useful, these studies have many inherent limitations (for review see^9^). One pivotal drawback is determining whether human post-mortem findings are specific to a trait or confounded by co-occurring illness, various technical and/or biological biases. Additional complexities exist among drug use disorder research as it is challenging to separate the effects of substance exposure history, drug toxicity and underlying mechanism of addiction. Synchronizing gene expression from precarious post-mortem human data with precise and carefully controlled studies in animal models affords the opportunity to pinpoint and characterize certain neuro-molecular sources of variability; it may also help quantify the pathogenicity of a specific behavior with a diseased state. Such insights will also inform the ongoing dialogue surrounding comparative cross-species approaches in bio-behavioral research^10^.

The current study queried extant publically available resources to identify transcriptome-wide studies of animal cocaine self-administration and human cocaine use disorder (CUD; DSM-IV abuse and/or dependence). We utilized data from a recent cocaine use/self-administration study that assessed the transcriptomic alterations in multiple reward circuitry regions in mice^11^. We compared this with two post-mortem studies of individuals diagnosed with CUD versus cocaine-free controls with analogous brain regions – the hippocampus^12^ and the midbrain/ventral tegmental area (VTA^13^) Previous cross-species gene expression studies for substance use have been limited to a single brain region and typically have relied on a limited focus on candidate targets or systems^14–16^. Given the etiology of substance use disorders encompasses many neurological substrates and involves a multitude of molecular systems, research is warranted to examine various tissues and assess global convergence across the transcriptome. Additionally, genes/transcripts tend to act in a broader biological context (e.g., gene co-expression networks), highlighting the importance of investigating cross-species convergence for individual genes and interconnected systems of genes.

We hypothesized that, within meso-limbic reward circuitry 1) individual genes associated with mouse cocaine self-administration would moderately correspond to homologous human genes/transcripts correlated with human CUD and that 2) systems (or networks) of co-expressed genes would demonstrate appreciable transcriptional convergence across mice and men.

## Materials and Methods

### Human Samples

We obtained male post-mortem human hippocampal (CA4 to CA1 and dentate gyrus) RNA-sequencing (RNA-seq) data from the Sequence-Read-Archive (accession#: SRA029279). After outlier screening^16^, this sample included seven samples with a diagnosis of CUD and eight drug-free controls (*M*_age_=39.3, *s.d.*=6.2, range=30-50; 66.67% European American; 33.33% African American). Samples from patients with CUD displayed a chronic pattern of cocaine use before death, died from cocaine toxicity and had prior diagnoses of cocaine abuse or dependence^12^ (DSM-IV). Note, since DSM-IV cocaine abuse and dependence have been reclassified into a singular DSM-V disorder, we defined these traits as CUD. Control subjects had no history of drug use and died from sudden cardiac or accidental events. All subjects were carefully matched on post-mortem interval (PMI; hours from death to freezer), ethnicity and age. No significant differences were observed for age and PMI across cocaine users and controls |*t*| < 0.51, all *p* > 0.623.

Leveraging publically available microarray data from the Gene Expression Omnibus (accession #: GSE54839), we used human post-mortem midbrain samples from twenty males^13^ (*M*_age_=49.2, *s.d.*=3.9, range=45-59; 80.0% African American, 20.0% European American). Each individual had three technical replicates, such that, a single person’s sample was run on the HT-12 Bead Chip microarray platform three separate times. Extraction of midbrain samples, which included the ventral tegmental area (VTA) and the substantia nigra (SN), was guided by the presence of neuromelanin and demonstrated enrichment for dopamine transcripts (Bannon et al. 2014). Half of the samples had CUD (a documented history of DSM-IV cocaine abuse) and died from cocaine abuse, toxicity or cocaine-related aortic complications. The other half of midbrain samples had no history of drug or cocaine abuse and died from acute causes such as cardiovascular disease or gunshot wounds. Cases and controls were well matched on race and did not differ on RNA integrity (degree of RNA degradation), age and brain pH level, all |*t*| < 1.04, all *p* > 0.313.

### Mouse Samples

We utilized classic cocaine-self administration RNA-seq data from male mice in two brain regions: the ventral hippocampus (n=15), and the VTA^11^ (n=12; Gene Expression Omnibus accession #: GSE110344). Note this sample consisted of genetically homogenous C57BL/6J mice (6-8 weeks old) that had *ad libitum* access to food and water, but restricted access to these resources during training and testing. After standard food training, jugular vein catheterization surgery was performed and mice recovered for 3-5 days before starting self-administration. Mice were randomly assigned to either press a lever for cocaine (0.5 mg/kg/infusion; 20 second time-out; n=6-8 per group/brain-region) or to press a lever for saline (n=6-7 per group/brain-region). Over 10-15 days mice underwent 2-hour self-administration sessions on either a fixed-ratio one (days 5-10) or a fixed-ratio two schedule (for 4-5 days). Twenty-four hours after their last self-administration session mice were euthanized.

In addition to classic cocaine-self administration, the same study used a *separate/independent cohort* of male C57BL/6J mice to evaluate other cocaine-related behaviors (see^11^ for more details). Briefly, this cohort experienced classic cocaine or saline self-administration, but after their last trial they experienced a 30-day forced abstinence period before being put in the following test conditions: 1) acute cocaine or saline exposure (among saline self-administering mice; 10 mg/kg intraperitoneal injection of cocaine), 2) re-exposure to only the *context* of cocaine or saline self-administration (context re-exposure) and 3) re-exposure to cocaine or saline in the context of cocaine/saline self-administration (cocaine re-exposure; 10 mg/kg intraperitoneal injection of cocaine). After removal of two outliers (due to low read counts), we used a total of 10-12, 9-11 and 11 mouse hippocampus and VTA samples for acute cocaine exposure, context re-exposure and cocaine re-exposure groups, respectively. To control for age differences, these mice were euthanized at the same age as the other cohort of mice and all mice experienced cocaine self-administration post-puberty. This independent cohort was primarily used to test the validity of gene networks constructed from the classic cocaine self-administration mice and was also leveraged for *post-hoc* comparative analyses (see Supplementary Information).

### Data Preparation

The current study processed RNA-seq with a unified approach. Specifically, we used Trimmomattic^17^ to remove Illumina adapters and low quality reads (Phred score < 12). Then, utilizing the Spliced Transcripts Alignment to a Reference tool^18^ we aligned RNA-seq data to either the human hg19 genome (*Homo sapiens*) or the mouse mm10 genome (*Mus musculus*). Our RNA-seq read alignment exceeded recommended thresholds19 for human (*M*_%_reads_aligned_=70.60%) and mouse RNA-seq data (*M*_%_reads_aligned_ =91.89%). In total, we mapped an average of 13,623,098 reads (*s.d*.=2,889,467) to the human hg 19 genome and 26,504,017 reads (*s.d*.=6,8965,611) to the mouse mm10 genome. Lastly, we transferred mapped RNA-seq reads into discrete RNA-seq counts via HTseq_0.6.1^20^ for bioinformatics analyses.

Microarray data from the human midbrain/VTA had three technical replicates per sample/person. Correlations across technical replicates were strong (all Pearson’s *r* > 0.95) suggesting that gene expression could be collapsed across eplicates. Therefore, we averaged microarray feature intensity recordings for individual array ids across technical replicates for each person/individual. Occasionally, multiple array ids exist for one individual gene, so we distilled array ids into a single gene-level expression value via the *collapseRow* function in R.

To account for gene expression variability among individual datasets, species and brain regions, we normalized data for each specific dataset separately (e.g., within species and within region). Performing normalization of only overlapping genes across species is thought to bias down-stream analyses^21^. Hence, we normalized transcriptome-wide data using all measured genes specific to each species and brain region, and consequently merged human and mouse datasets to homologous genes listed from the Mouse Genome Informatics website (http://www.informatics.jax.org/). Improperly accounting for differences across RNA-seq and microarray platforms can also be a source of bias for bioinformatic results, thus we harmonized our gene expression normalization using the *voom (RNA-seq)* and *vooma (microarray*) commands.

### Analyses

To identify individual genes associated with both cocaine use/self-administration and CUD, we compared associations across human and mouse differential expression analyses. These analyses sought to minimize potential biases from analytical techniques by using the same differential expression platform (*limma*) that conducted empirical Bayes models^22^. To control for technical noise, all differential expression analyses adjusted for hidden batch effects using the svaseq package, which stabilizes error estimates^23^. Log fold change estimates from differential expression analyses are still estimated with noise – especially for genes with low baseline expression. To account for this noise, we focused our analyses on differential expression *t*-statistics, which balance effect size and standard errors for log fold change estimates, and thus, enabled us to (more confidently) characterize comparative gene expression signatures transcriptome-wide. We quantified the association of cross-species gene expression associations using standard linear regressions and Pearson product-moment correlations. Cross-species differential expression analyses included a range of 10,869 to 12,721 homologous genes/transcripts. Differentially expressed genes that demonstrated associations in the same direction with mouse cocaine self-administration and human CUD were required to surpass a differential expression cutoff of |*t*| > 2. We annotated these differentially expressed genes with the Kyoto Encyclopedia of Genes and Genomes (KEGG; Mouse 2019) pathways using Enrichr^24^. To control for false positives, we required significant KEGG pathways to survive correction for multiple testing (*p*_adj_ < 0.05) and surpass an enrichment odd’s ratio (OR) > 2.00.

Next, we performed cross-species comparisons of gene *co-expression* patterns. First, we created signed WGCNA gene co-expression networks^25^ from mouse cocaine self-administration data. For computational limitations, we filtered out lowly expressed genes/transcripts (< 0.1 read count per sample), which resulted in 16,029 and 15,751 genes/transcripts for WGCNA modeling in the hippocampus and VTA, respectively. Using these genes/transcripts, we created a correlation matrix of the expression of all WGCNA genes/transcripts with themselves. We weighted these co-expression matrices by raising the correlations to a power of 12 in the hippocampus and a power of 7 in the VTA, which maximized WGCNA model fit (all scale free topologies > 0.86). Our study transformed weighted correlation matrices into topological overlap matrices using the *TOMdist* command, which we converted into distance metrics (Euclidean distance) and then further split into clusters of correlated genes using the *hclust* command. To make discrete WGCNA gene networks we parsed clusters of correlated genes via a standard dynamic tree-cutting algorithm (minimum network size = 50). In total, our WGCNA modeling created 22 hippocampal and 19 VTA gene co-expression networks, which were arbitrarily assigned to a color.

To test the stability, validity and *conservation* of these WGCNA networks, we used a three-step network preservation procedure similar to Vanderlinden et al.^26^. WGCNA modeling in small sample sizes may yield unstable gene networks (e.g., created by chance). Therefore, to test whether mouse WGNCA networks were stable and of high quality, we used a within sample technique to assess network reproducibility/preservation. Specifically, we assessed the degree of network preservation of WGCNA networks by comparing them to 100 bootstrapped samples from the original datasets (stability). Then we quantified the validity of these WGCNA gene networks. We defined validity as whether WGCNA gene networks could be re-created in the same tissue and species *but* in an independent cohort (of cocaine or saline exposed mice). Lastly, we tested the *conservation* of WGCNA gene networks within tissue and across species. If mouse WGCNA gene networks were conserved, it would suggest that co-expression patterns from self-administering mice were reproducible in disordered human cocaine users and controls. As recommended by Langfelder et al.^27^, all WGCNA network preservation analyses used *Z*_summary_ statistics, which aggregate across seven measures of gene network reproducibility. *Z*_summary_ statistics below a score of 2 indicate that networks/co-expression patterns are not reproducible, whereas scores between 2-10 indicate weak to moderate reproducibility and *Z*_summary_ statistics above 10 are thought to be highly robust^27^.

Since hub genes are thought to represent biologically meaningful targets in disease progression^28^, we focused our analyses on hub genes from the most conserved gene networks. We defined hub genes as the top 1% of most intra-connected network genes (e.g., the most co-expressed with all other genes in the network). To better characterize the functional role of these genes in the brain we performed expression-based phenome-wide association analyses (ePheWASs^29^; https://systems-genetics.org/). Specifically, these analyses investigated the associations between the expression of a hub gene, in a specific brain region, with 1,250 unique “nervous system” traits – including both molecular and behavioral phenotypes - from the BXD mouse population (consisting of inbred lines derived from the progeny of a cross between C57BL/6J and DBA/2J mice). Significant ePheWAS were required to survive a Bonferroni *p*-value correction for multiple testing (*p*=0.05/1,250 or 0.0004).

## Results

### Differential Expression – Individual Genes

We conducted differential expression analyses within species and tissues to identify specific cocaine-related genes and then quantified the degree of overlap between mice and humans. The genes with increased expression in cocaine self-administering mice tended to also be increased in humans with CUD, in both the hippocampus*, b*=0.076, *s.e.*=0.014, *F*_(1, 12719)_=29.82, *p*=4.823e-8, *R*2=0.0023, and VTA/midbrain, *b*=0.042, *s.e*=0.001, *F*_(1, 11856)_=17.83, *p*=2.432e-5, *R*2=0.0014 (see **Fig 1**). While these associations were significant, in the expected direction and reproducible across brain regions, they were modest in magnitude and only accounted for ~0.1%-0.2% of the molecular variance observed in human CUD.

**Figure 1.**
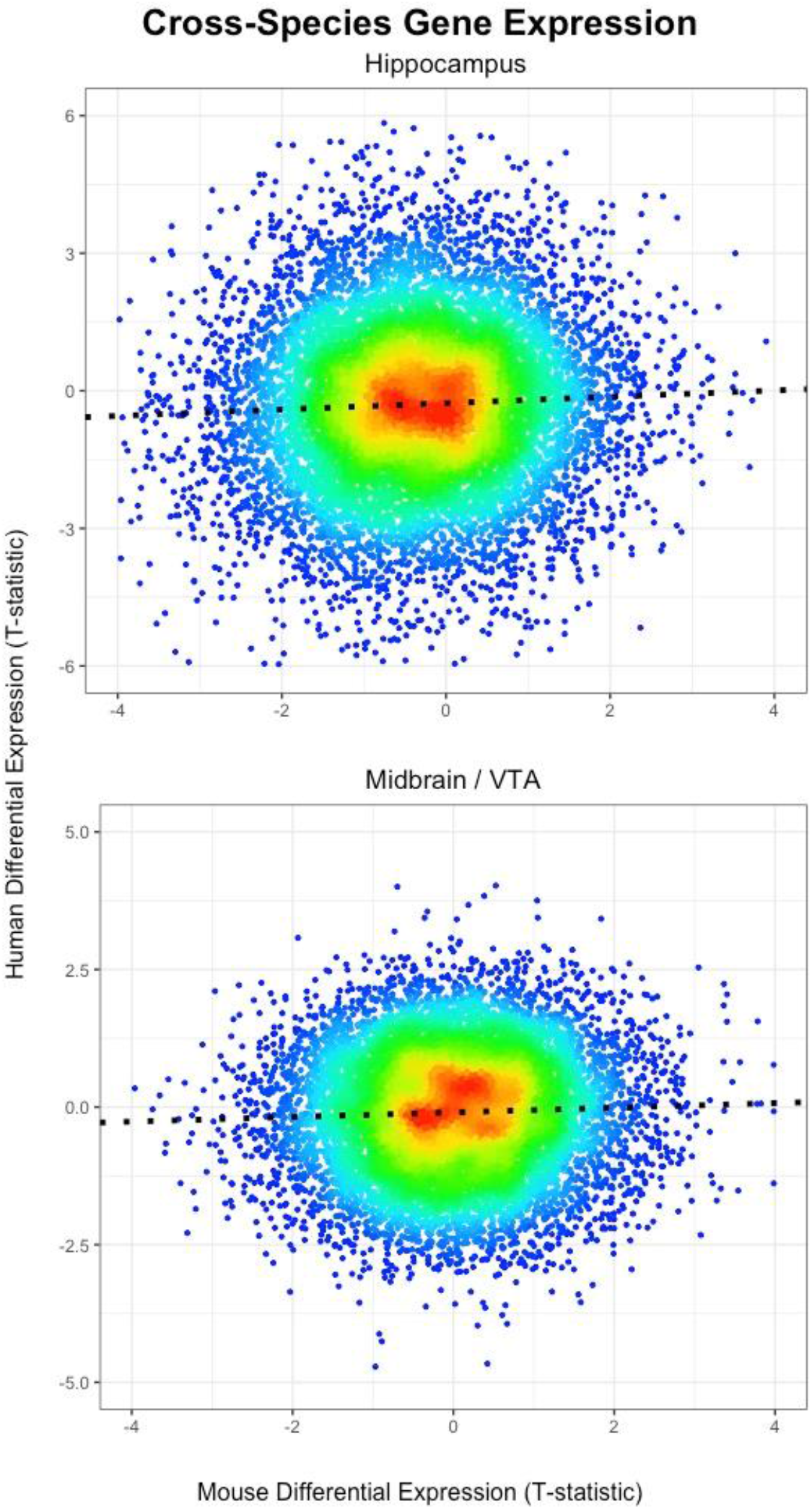
Correlating Differentially Expressed Genes across Mouse Cocaine Self-Administration and Human CUD Heat scatter plots showing overlap of individual genes/transcripts associated with cocaine self-administration in mice (x-axis; n=12-15) and human CUD (y-axis; n=15-20) in the hippocampus (top) and midbrain/VTA (bottom). Note that each dot represents a gene/transcript, which are color coded by their density, with high frequency/density regions shown in red and low frequency genes/transcripts represented in blue.

To characterize genes bolstered with cross-species support, we performed functional annotation analyses of the differentially expressed genes analogously associated with cocaine self-administration and CUD. Collapsing across brain regions and using a *t*-statistic cutoff of > |2|, we found a total of 173 genes that were associated with both mouse cocaine self-administration and human CUD in the same direction (see Supplementary File S1). These genes were enriched for thermogenesis, neurodegenerative diseases, glycolysis/gluconeogenesis, oxidative phosphorylation and neurotrophin signaling (see Supplementary Table 1). Together, these results quantified the extent of transcriptional conservation and disentangled the biological targets underlying mouse cocaine use and human CUD. We also performed *post-hoc* analyses that evaluated the cross-species differential expression overlap from three additional cocaine behaviors in mice: 1) acute cocaine exposure, 2) context re-exposure and 3) cocaine re-exposure. But, we found no replicable associations with human CUD across brain regions (see Supplementary Figure S1).

### Gene Co-expression Analyses – Systems of Genes

Our next step was to address the cross-species correspondence of biological systems in the hippocampus and VTA/midbrain. Using mice from the classic cocaine self-administration model, we created 22 data-driven WGCNA gene co-expression networks in the hippocampus and 19 WGCNA gene co-expression networks in the VTA. We focused our analyses on the stable (reproducible within sample) and valid (preserved in an independent mouse cohort) gene co-expression networks, which included 14-gene co-expression networks in the hippocampus (all Z_summary_ > 2.017) and eight in the VTA (all Z_summary_ > 3.367; see **Fig 2**). We investigated whether these robust WGCNA networks (gene co-expression patterns) were *conserved* within-tissue and across species. In the hippocampus, only the turquoise WGCNA gene network (3,218 genes) demonstrated moderate conservation in human CUD and controls, Z_summary_=8.793. In the mouse VTA, two gene co-expression networks demonstrated weak to moderate conservation in the human midbrain: the green-yellow network (212 genes; Z_summary>_=2.631) and the blue network (3,726 genes; Z_summary_=6.168).

**Figure 2.**
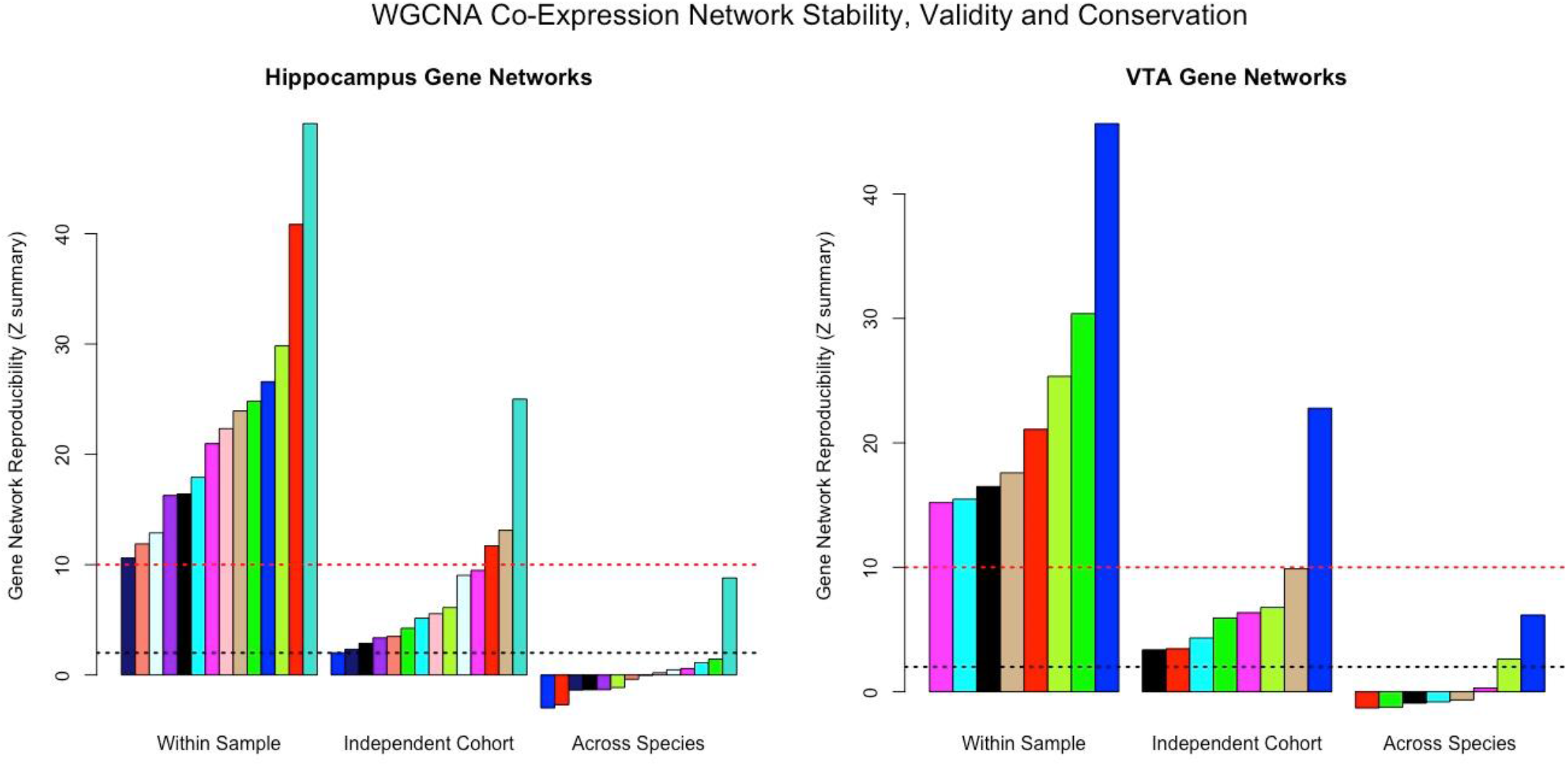
Reproducibility of Gene Co-expression Networks Barplot displaying the reproducibility/robustness of WGCNA gene co-expression networks derived from the mouse hippocampus (left) and VTA (right). Each bar represents a specific gene network - each arbitrarily assigned to a color - and constructed from cocaine self-administration mouse data. The three sections of the x-axes show whether gene networks were reproducible within sample (100 bootstrapped samples), in a separate group of cocaine/saline-exposed mice (independent cohort/same study; Walker et al. 2018) *or* conserved in post-mortem human CUD/controls (across species). The y-axes are the Z_*summary*_ preservation statistic values. Bars surpassing the dashed red line indicate gene networks that are highly robust and bars that exceed the dashed black line are considered weak to moderately reproducible. *Note* the turquoise hippocampal gene network and blue VTA gene network are highly stable/valid and are weak to moderately conserved in human tissue

We selected the most conserved WGCNA gene co-expression networks per brain region for follow-up investigation - the turquoise hippocampal network (Supplementary File S2) and blue VTA network (Supplementary File S3). These data-driven gene co-expression networks were biologically meaningful and demonstrated enrichment for both shared and specific KEGG pathways including: cocaine/drug addiction, neurotransmitter signaling, synaptic plasticity processes and various other KEGG pathways (see **Table 1**). We further investigated the overlap between the turquoise and blue gene networks and found over 2,000- shared genes across networks (see Supplementary Figure S2). Despite, the substantial similarities in gene co-expression network structure, we found minimal overlap of differentially expressed genes (*p* < 0.05) across the hippocampus and VTA from the mouse cocaine self-administration paradigm (see Supplementary Figure S3). Overall, these results suggest substantial transcriptional overlap across brain regions at the systems level but not at the individual gene/transcript level.

**Table 1.** KEGG Pathway Enrichment for Conserved WGCNA Gene Co-expression Networks This table shows significantly enriched KEGG pathways (*p*_adj_ < 0.05 & OR > 2.0) common and specific to the turquoise hippocampal and blue VTA gene co-expression networks. *Note* we highlighted in grey that the blue VTA gene network was enriched for the Cocaine Addiction KEGG pathway.

| Potential Functions of Conserved Systems/Networks Across Humans and Mice |  |  |  |  |  |  |
| --- | --- | --- | --- | --- | --- | --- |
| KEGG Pathway | Hippocampus (Turquoise) |  |  | VTA (blue) |  |  |
|  | OR | *p*-value | *p*<sub>adj</sub> | OR | *p*-value | *p*<sub>adj</sub> |
| Synaptic Vesicle Cycle | 3.25 | 3.19E-13 | 3.12E-11 | 3.04 | 1.87E-13 | 2.83E-11 |
| Alzheimer's Disease | 2.46 | 6.92E-14 | 1.05E-11 | 2.18 | 1.63E-11 | 1.23E-09 |
| Oxidative Phosphorylation | 2.47 | 4.62E-11 | 3.50E-09 | 2.16 | 7.23E-09 | 2.19E-07 |
| Parkinson's Disease | 2.30 | 1.03E-09 | 4.46E-08 | 2.31 | 1.90E-11 | 1.15E-09 |
| Long Term Potentiation | 2.70 | 1.09E-07 | 2.53E-06 | 2.60 | 3.70E-08 | 9.33E-07 |
| Dopaminergic Synapse | 2.13 | 1.87E-07 | 3.15E-06 | 2.22 | 1.05E-09 | 4.56E-08 |
| GABAergic synapse | 2.36 | 4.79E-07 | 6.30E-06 | 2.24 | 4.14E-07 | 6.60E-06 |
| Glutamatergic synapse | 2.30 | 5.23E-08 | 1.32E-06 | 2.10 | 3.02E-07 | 5.72E-06 |
| Cardiac muscle Contraction | 2.16 | 4.40E-05 | 3.10E-04 | 2.10 | 2.39E-05 | 2.69E-04 |
| Endocytosis | NS | NS | NS | 2.15 | 3.64E-16 | 1.10E-13 |
| Amphetamine Addiction | NS | NS | NS | 2.57 | 5.76E-08 | 1.34E-06 |
| Ubiquitin Mediated Proteolysis | NS | NS | NS | 2.02 | 1.82E-07 | 3.94E-06 |
| Nicotine Addiction | NS | NS | NS | 2.45 | 9.55E-05 | 7.42E-04 |
| *Cocaine Addiction* | NS | NS | NS | 2.27 | 1.48E-04 | 9.95E-04 |
| Long Term Depression | NS | NS | NS | 2.06 | 2.91E-04 | 1.80E-03 |
| ... Sodium Reabsorption | NS | NS | NS | 2.01 | 5.51E-03 | 2.20E-02 |
| Axon Guidance | 2.15 | 2.96E-10 | 1.79E-08 | NS | NS | NS |
| ... Endocannabinoid Signaling | 2.08 | 1.23E-07 | 2.67E-06 | NS | NS | NS |
| Adrenergic Signaling .... | 2.07 | 2.08E-07 | 3.31E-06 | NS | NS | NS |
| Spliceosome | 2.13 | 2.49E-07 | 3.76E-06 | NS | NS | NS |
| ErbB Signaling Pathway | 2.23 | 8.36E-06 | 7.45E-05 | NS | NS | NS |
| Gap Junction | 2.18 | 1.43E-05 | 1.14E-04 | NS | NS | NS |
| SNARE ... in Vesicular Transport | 2.27 | 3.63E-03 | 1.36E-02 | NS | NS | NS |
| Circadian Rhythm | 2.08 | 1.50E-02 | 4.46E-02 | NS | NS | NS |

We concentrated on hub genes for the moderately conserved turquoise and blue gene co-expression networks. In total, we identified 32 and 37 hub genes for the turquoise (hippocampus) and blue (VTA) gene networks, respectively. Note that *nine hub genes* were in the top 99% of gene network intra-connectedness in *both* the turquoise and blue gene co-expression networks, which we defined as common hub genes. We visualized both the common and specific hub gene structure across networks/brain regions (see **Fig 3**). Our results suggest that common hub genes could have an important role in multiple brain regions, but the function of these hub genes in the context of cocaine use is not fully understood.

**Figure 3.**
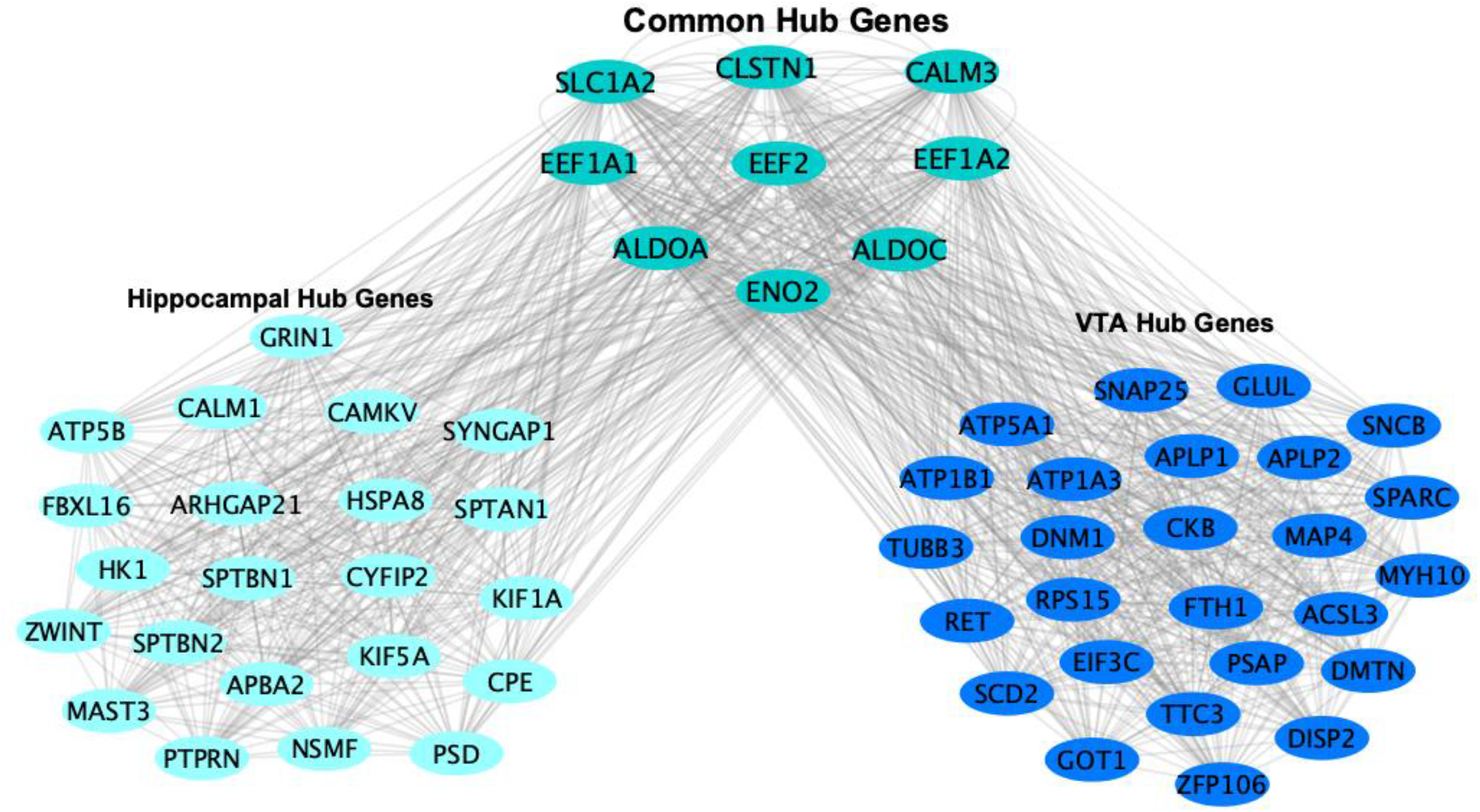
Common and Specific Architecture of Hippocampal and VTA Gene Networks – Hub Genes Hub genes are defined as being in the top 99^th^ %tile of gene network intraconnectedness. Common hub genes are in the top 1% of gene network connectedness in *both* the turquoise hippocampal gene network (left) and blue VTA gene network (right). Region specific hub genes are in the top 1% of their respective gene network but not in the top 1% of connectedness in the other gene network.

To investigate specific behavioral mechanisms related to the expression of our common hub genes in mice, we conducted ePheWASs - first in the VTA and hippocampus and subsequently in eight other brain regions, including: the amygdala, cerebellum, hypothalamus, nucleus accumbens, striatum, pituitary, prefrontal cortex and whole brain. In total, our common hub genes exhibited eighteen significant neuro-transcriptional associations (with thirteen unique mouse traits) in eight different brain regions (see **Fig 4**). These results suggest that our common hub genes exert a broad functional role spread across the brain (see Supplementary File S4). Most significant ePheWAS associations were molecular. *Slc1a2, Calm3, Aldoa, Aldoc* and *Eno2* were associated with dopamine and serotonin function in the VTA and striatum. Additionally, we found significant ePheWAS associations between *Eef1a1, Aldoa* and *Eef1a2* with alcohol conditioned place aversion^30^ cocaine conditioned place preference^31^ and cocaine self-administration^32^, respectively. Overall, these analyses better characterize the behavioral and neuro-molecular role of our common hub genes within particular brain regions.

**Figure 4.**
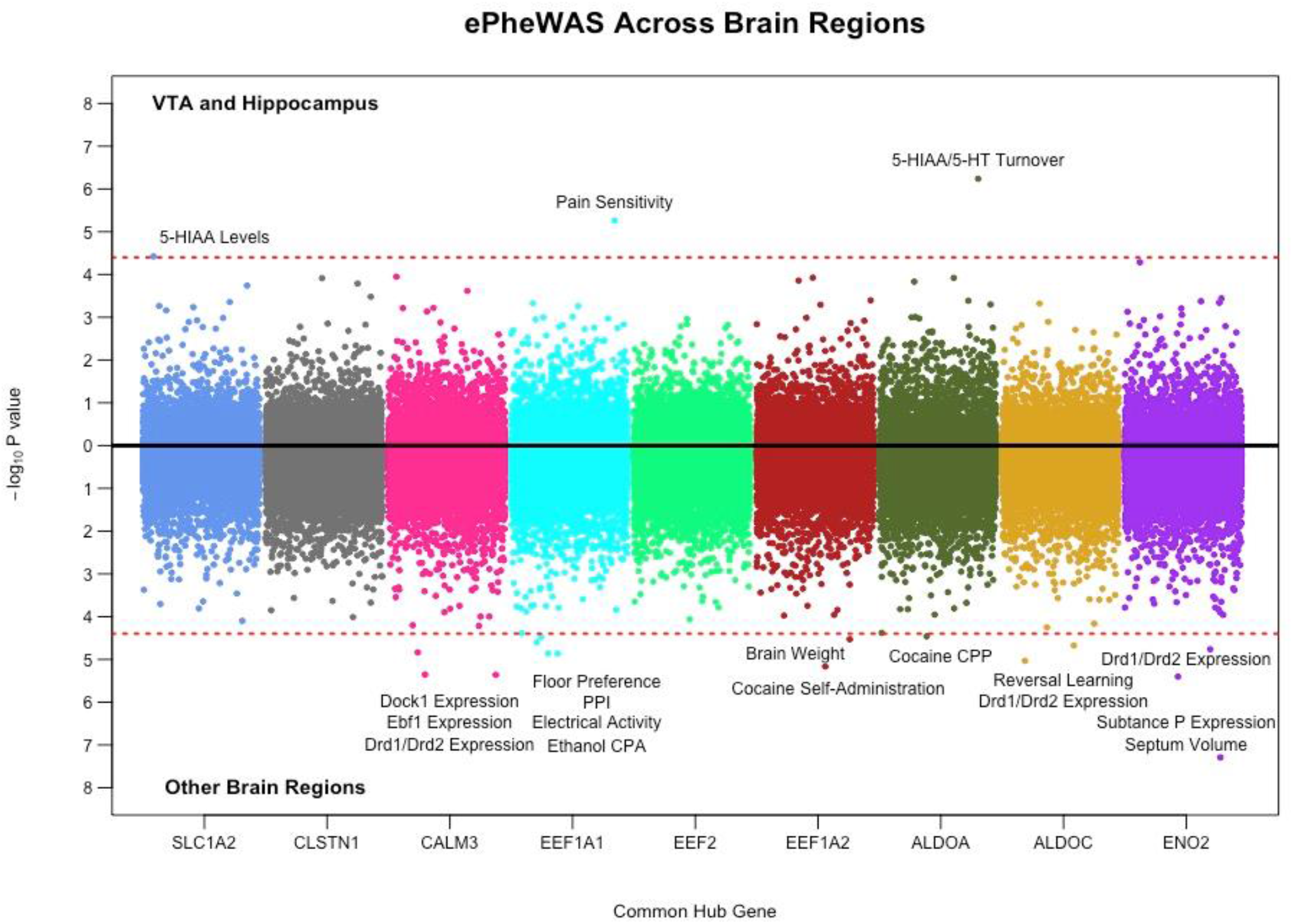
Common Hub Genes Expression-based Phenome-wide Association Study (ePheWAS) of Nervous System Traits in Mice Miami plot showing the results of our ePheWASs of common hub genes from the turquoise and blue gene networks. Each dot is a nervous system phenotype/trait (1,250 unique traits), which are color coded by hub gene (x-axis). The y-axis represents the −log_10_ p-value from the association of a hub gene’s expression with a particular trait in a certain brain region. The top part of the plot shows results from the VTA or hippocampus and the bottom includes results from eight other cortical/sub-cortical brain regions. The dashed red line denotes the Bonferonni correction for multiple testing (0.05/1,250) and thus, all dots surpassing this threshold indicate significant associations between the expression of a gene with a trait. *Note* that our ePheWAS analyses utilized BXD mice, which are derived by crossing C57BL/6J and DBA/2J inbred mice. PPI = pre-pulse inhibition; CPP = conditioned place preference; CPA = conditioned place/taste aversion.

## Discussion

The current study identified modest to moderate conservation of transcriptional responses across mouse cocaine self-administration and human CUD within reward circuitry. Specifically, we found convergent gene expression for individual genes/transcripts *and* systems of co-expressed genes/transcripts in the hippocampus and midbrain/VTA for the ubiquitous short access, intravenous cocaine self-administration model. Our study is the first to quantify the extent to which meso-limbic gene expression findings from mouse models of cocaine self-administration directly recapitulate human CUD.

While the correlated neuro-transcriptional response between mouse cocaine self-administration and human CUD was significant, and in the expected direction, it was surprisingly modest. We considered various interpretations for these potentially sobering effect sizes. Inbred mice pressing a cocaine lever for two-weeks and individuals with CUD have different degrees of exposure duration, use severity and genetic backgrounds, which may recruit separate dimensions of biological variation. Perhaps the molecular mechanisms governing reward learning are akin to the development of substance use behaviors, but not necessarily tapping into the core pathogenic features of the addictive process, or at least the “end state.” Thus, the neuro-pathophysiology of compulsive human cocaine use may be far more complex than the transcriptional thumbprint of a well-controlled operant behavior. Alternative models of drug use may improve cross-species prediction. Although, acute cocaine exposure, context re-exposure and cocaine re-exposure models tended to be less robust predictors of human CUD than the classic self-administration paradigm (see Supplementary Figure S1).

On the other hand, human post-mortem brain tissue may be responsible for dampening cross-species effect sizes. If only a slim proportion of the observed human gene expression is specific to CUD then our cross-species effects may seem small but could be capturing a fair amount of disordered cocaine use variability and be obfuscated by potential human confounds (co-occurring illness, cocaine toxicity etc.). It is also possible that our cross-species comparisons do not follow a positive linear association. For instance, during drug tolerance, neurotransmission genes may demonstrate expression in one direction from short-term or acute use, but exhibit compensatory expression in the opposite direction through chronic drug exposure. In accordance with this, and previous cross-species research (Enoch et al. 2012), we found various receptor (*CHRNB2, GRIN1, GLYR1)* and transporter genes (*SLC12A3, SLC24A2, SLC35F5, SLC6A7*) differentially expressed in opposing directions between mouse cocaine self-administration and human CUD (all |t| > 2); see Supplementary File S5). The aforementioned interpretations are not mutually exclusive and, in practice, could act in a combinatorial way to mitigate cross-species associations. Regardless, the current study speaks to the difficulty of modeling and capturing the molecular relevance of specific animal systems for complex human diseases. We rely on models as simplifications of complicated traits, to enable examination of specific facets with precise experimental perturbations, but no model in isolation recapitulates the daunting complexity of the human experience.

We extend previous cross-species drug use research^33^ by investigating the convergence of *co-expression* patterns across species in the hippocampus and VTA/midbrain. While only a few of the mouse gene co-expression networks surpassed the threshold of (weak to moderate) reproducibility in analogous human tissues, these systems included thousands of genes. Reassuringly, we found that the most conserved gene co-expression networks included numerous candidates from the cocaine addiction pathway. To identify critical targets of these conserved systems, we honed in on the major players (hub genes), which may regulate the expression and/or function for a multitude of these genes. These genes tended to be involved in broad nervous system functions, such as: synaptic plasticity (*Cyfip2, Ret, Tubb3, Sptbn1, Sptbn2, Spatn2*), glutamate transmission (*Slc1a2, Grin1, Glul*) and calcium signaling (*Calm1, Calm2, Calm3, Atp1b1, Camkv*) potentially indicating a conserved and pleotropic role for more general dimensions of drug use and/or addiction. Accordingly, previous research demonstrates genome-wide associations of these genes with other human substance use disorder traits commonly co-morbid with cocaine abuse/dependence, including: alcohol (*Arhgap21^34^*), tobacco (*Kif1a^35^*), cannabis (*Ret^36^*) and heroin (*Myh10^37^*) dependence symptoms.

From the conserved VTA and hippocampus gene networks, we observed nine common hub genes that may modulate brain function in multiple regions. Notably, over half of these genes (*Calm3, Aldoc, Clstn1, Eef1a1 and Slc1a2*) were also hub genes in a PFC gene network associated with human CUD that was enriched for analogous KEGG pathways^38^. It is possible that common hub genes recruit similar biological systems relevant to cocaine addiction across various brain regions. The expression of our common hub genes were also associated with relevant traits from the mouse realm - such as dopamine and serotonin function (*Slc1a2, Calm3, Aldoa, Aldoc* and *Eno2*), cocaine reward and/or memory (*Aldoa*) and cocaine self-administration (*Eef1a2*) in the striatum, VTA, hypothalamus and cerebellum. In sum, we identify and characterize critical genes from conserved biological systems relevant to cocaine use and rank targets for follow-up investigation within particular tissues. Of note, 64.74% of differentially expressed genes analogously associated with cocaine self-administration and CUD were a part of the turquoise or blue gene networks, suggesting that these complementary individual-level and systems-level bioinformatics methods usurp both common and unique sources of transcriptional conservation.

The current study has various limitations. Ascertainment of brain tissues was performed in similar but not identical neurobiological substrates. That is, mouse tissues were collected from the ventral hippocampus and VTA, whereas human samples used tissue from the entire hippocampus and dopamine enriched regions of the midbrain (VTA and substantia nigra). Another inconsistency regarding our cross-species comparison is that human findings are a result of heterogeneous genetic predisposition and environmental exposure, whereas the C57BL/6J mice are genetically homogenous with very controlled environmental conditions. Additionally, while the current study utilized fairly small sample sizes, it is the largest comparative transcriptomic human study on cocaine use to date – including multiple brain regions, cocaine behaviors and analytical techniques.

In light of the current study, we suggest various directions for future research. Relatively large post-mortem human brain samples are publically available for alcohol^39^ and opiates^40^ and could be integrated with a multitude of traits from model systems. Animal models of compulsive drug use (e.g., escalated use, aversion resistant users etc.) have apparent face valid behavioral correspondence to clinical symptomology of drug use disorders^41^, but whether, or to what extent, these molecular mechanisms recapitulate genuine features of human addiction is not known. Because there are ten times more expression studies for cocaine use in animals than humans^42^, comparative neuro-transcriptomic work would benefit from selecting human post-mortem brain regions that are supported by the vast literature and bioinformatics database in animals (e.g., striatum/nucleus accumbens). Unfortunately, the largest post-mortem human brain study on cocaine was conducted in dorsal-lateral PFC neurons^43^, which seems to lack a homologous substrate in the rodent brain. Heterogeneity in cell types across species can pose an additional bias from cross-species comparisons^44^. Thus, a cleaner signal may emerge from integrating single cell bioinformatics analyses across species. Ideally, cross-species research will aggregate across various layers of the transcriptome and employ global integrative strategies incorporating a multitude of model systems and human traits to illuminate the molecular crossroads among particular models and disease processes.

In conclusion, this is one of the first studies to test the extent to which neuro-molecular correlates identified from animal models of cocaine use generalize to humans diagnosed with CUD. Our analyses characterize and provided insight into specific targets at the intersection of an established mouse paradigm and a corresponding human trait. The current study, thus, contributes to the burgeoning dialogue surrounding the pre-clinical, theoretical and empirical implications of animal research for human disease. We found that meso-limbic gene expression from a single, granular animal paradigm applied in a single strain of mice recapitulated only a modest fraction of the transcriptional effects observed in human disease. To fully capture the complexity of human addiction a broader spectrum of model organism diversity and experimental breadth needs to be encompassed in cross-species analyses.

## Acknowledgements

We express our gratitude towards the funding sources that supported our project: P50 DA039841, R01 DA037927 and DP1 DA042103.

## Author contribution

Spencer B. Huggett performed all analyses and wrote the initial manuscript. Rohan H. Palmer, Jason A. Bubier and Elissa J. Chesler aided in revising the manuscript, interpreting the findings and offered insight into what analyses would be appropriate for cross-species analyses

