## Supplementary_Information for "Meso-limbic Gene Expression Findings from Mouse Cocaine Self-Administration Recapitulate Human Cocaine Use Disorder": Cross_Species_Supplement.docx

*Post-hoc Analyses*

We examined whether alternative animal models may better explain the molecular signatures of human cocaine use disorder using an independent cohort of mice (from the same mouse study; Walker et al. 2018) that evaluated three different cocaine behaviors: 1) acute cocaine exposure, 2) context re-exposure and 3) cocaine re-exposure. Controlling for batch effects, we performed differential expression analyses for each individual behavior and brain region and then assessed the correlations of the resultant differential expression *t*-statistics across all traits. We identified significant hippocampal associations between acute cocaine exposure, context re-exposure and cocaine re-exposure with human cocaine use disorder – although some were estimated in the opposite direction than expected. In the VTA/midbrain, we found no significant cross-species associations of these behaviors (see Supplementary Figure S1).

**Supplementary Table S1** Significant KEGG Pathway Enrichment for Individual Genes Associated with Mouse Cocaine Self-Administration and Human Cocaine Use Disorder via Differential Expression Analyses

| **Function of Similarly Differentially Expressed Genes Across Species** | | | |
| --- | --- | --- | --- |
| Term | OR | *p*-value | *p*_adj_ |
| Thermogenesis | 5.00 | 3.42E-05 | 0.0104 |
| Alzheimer's Disease | 5.28 | 1.45E-04 | 0.0147 |
| Glycolysis / Gluconeogenesis | 8.63 | 2.86E-04 | 0.0173 |
| Non-alcoholic Fatty Liver Disease | 5.36 | 3.48E-04 | 0.0176 |
| Parkinson's Disease | 5.62 | 2.61E-04 | 0.0198 |
| Propanoate Metabolism | 14.92 | 1.42E-04 | 0.0215 |
| Neurotrophin Signaling Pathway | 5.73 | 6.49E-04 | 0.0281 |
| Oxidative Phosphorylation | 5.18 | 1.11E-03 | 0.042 |

To increase power for KEGG enrichment analyses, we collapsed similarly expressed genes across brain regions. We only included genes that were expressed in the same direction across traits/species and yielded a t-value > |2| (see Supplementary File S1).

**Supplementary Figure S1** Post-hoc Analyses: Gene Expression Correlations Across Mouse Models of Cocaine use and Human Cocaine Use Disorder

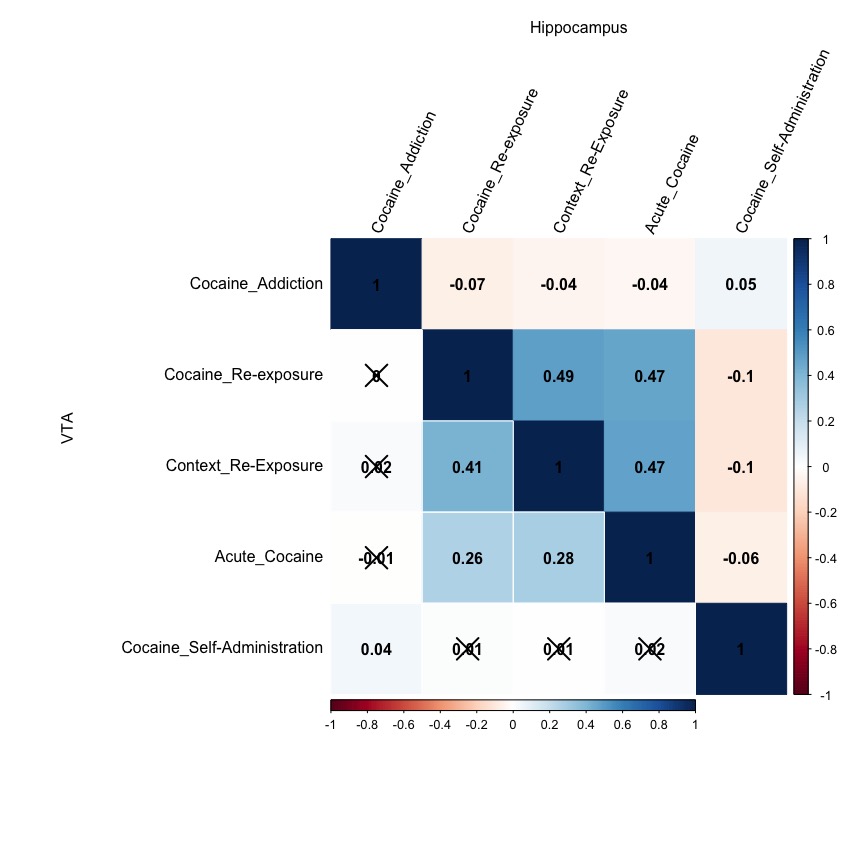

f

Transcriptome-wide correlation matrix of differential expression results (*t-*statistics) across specific cocaine behaviors in mice and human cocaine addiction (cocaine use disorder) within the hippocampus (upper diagonal) and VTA/midbrain (lower diagonal). Pearson product correlation coefficients are shown in black text and those that did not survive a Bonferroni correction for multiple testing (*p* < 0.05/10) were crossed out. Correlation strength was color coded with blue representing positive associations and blue displaying negative correlations.

**Supplementary Figure S2** Venn Diagram Demonstrating the WGCNA Gene Co-expression Network Structure Across the Mouse Hippocampus and VTA

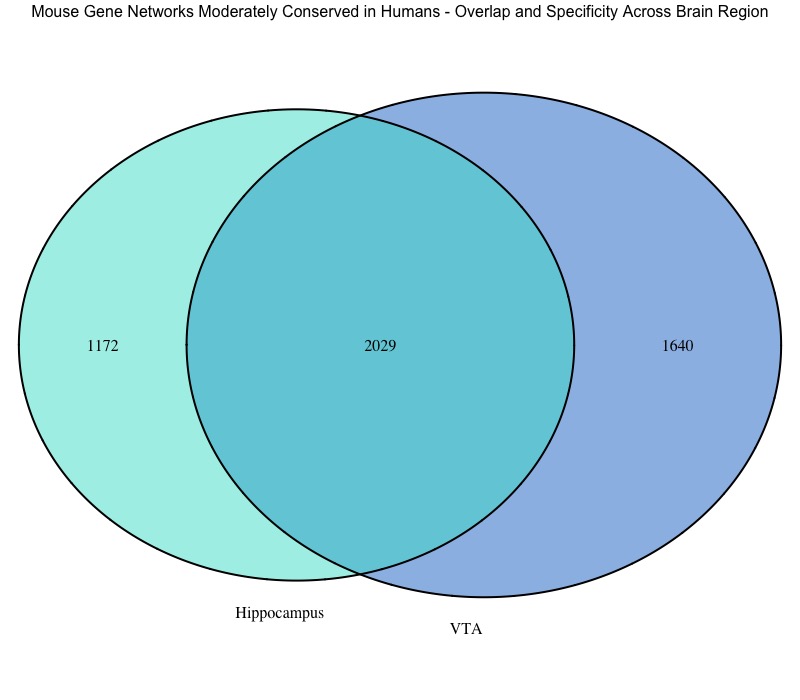

Venn diagram is color coordinated to match the WGCNA co-expression networks with the left circle showing the # of genes from the turquoise hippocampal network and the right circle representing the blue VTA gene network. Note that the observed overlap for the # of shared genes across gene networks/brain region is more than we would expect due to chance, OR = 10.17, 95% CI [9.30, 11.13], p < 2..2e-16, as tested via a Fisher’s exact test or Jaccard Similarity = 0.42, *p* < 0.002 (see Baker et al. 2013).

**Supplementary Figure S3** Venn Diagram Demonstrating the of Overlap/Specificity of Individual Genes Associated with Cocaine Self-administration Across the Mouse Hippocampus and VTA

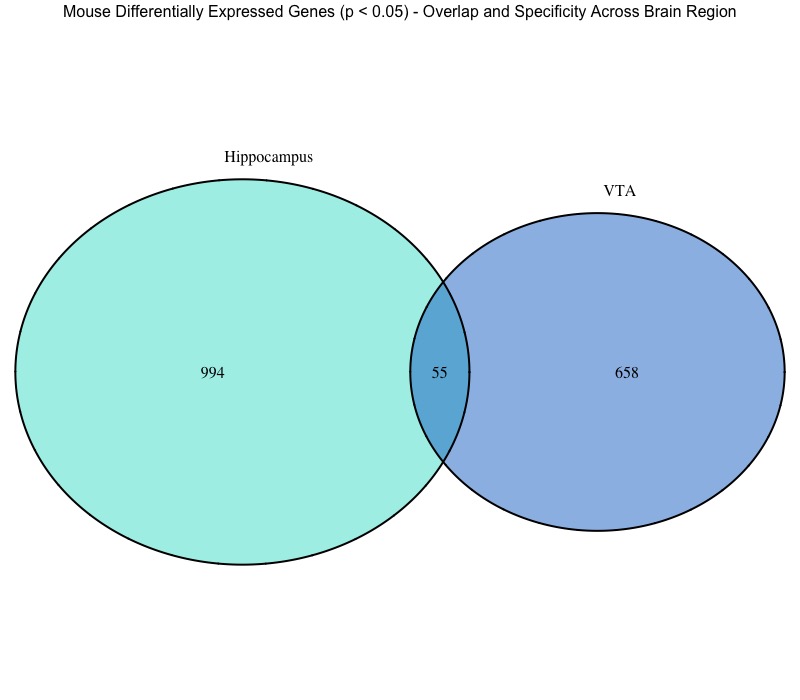

This venn diagram shows the number of differentially expressed genes (*p* < 0.05) associated with mouse cocaine self-administration in the hippocampus (left) and the VTA (right). Note that the observed overlap for the # of shared individual genes from differential expression analyses across brain regions is not more than we would expect due to chance, OR = 1.07, 95% CI [0.79, 1.43], *p* > 0.05, as tested via a Fisher’s exact test and Jaccard Similarity = 0.00, p > 0.05 (see Baker et al. 2012). Additionally, we found no significant linear association between t-statistics between hippocampal and VTA differential expression analyses, *β* = 0.015, *s.e.* = 0.008, *p* > 0.05, *R*^2^ =0.0002, via a standard linear regression

*References*

Baker, E. J., Jay, J. J., Bubier, J. A., Langston, M. A., & Chesler, E. J. (2012). GeneWeaver : a web-based system for integrative functional genomics. *Nucleic Acids Research*, *40*, 1067–1076. https://doi.org/10.1093/nar/gkr968
